# ASSESSMENT OF STABILIZING FEEDBACK CONTROL OF WALKING, A TUTORIAL

**DOI:** 10.1101/2023.09.28.559516

**Authors:** Jaap van Dieën, Sjoerd Bruijn, Maarten Afschrift

## Abstract

Walking without falling requires stabilization of the trajectory of the body center of mass relative to the base of support. Model studies suggest that this requires active, feedback control, i.e., the nervous system must process sensory information on the state of the body to generate descending motor commands to the muscles to stabilize walking, especially in the mediolateral direction. Stabilization of bipedal gait is challenging and can be impaired in older and diseased individuals. In this tutorial, we illustrate how gait analysis can be used to assess the stabilizing feedback control of gait. We present methods ranging from those that require limited input data (e.g. position data of markers placed on the feet and pelvis only) to those that require full-body kinematics and electromyography. Analyses range from simple kinematics analyses to inverse dynamics. These methods assess stabilizing feedback control of human walking at three levels: 1) the level of center of mass movement and horizontal ground reaction forces, 2) the level of center of mass movement and foot placement and 3) the level of center of mass movement and the joint moments or muscle activity. We show how these can be calculated and provide a GitHub repository (https://github.com/VU-HMS/Tutorial-stabilizing-walking) which contains open access Matlab and Python code to calculate these. Finally, we discuss what information on feedback control can be learned from each of these.

## 1. INTRODUCTION

Preventing a fall while walking requires stabilization of the position of the body center of mass over the base of support. The common occurrence of falls in toddlers who learn to walk, illustrates the difficulty of this control process. Falls are quite common also in older adults and in people with orthopedic or neurological disorders (Lord et al., 2001; WHO, 2007) and more importantly in these groups the consequences of falls are often severe (MacIntyre & Dewan, 2016; Talbot et al., 2005; Wenning et al., 1999).

From a mechanical perspective, it is not surprising that falls are common during walking. The mechanical challenge in human bipedal gait arises from our bodies being upright and segmented and our small base of support, particularly in single stance. Additionally, the anatomy is “top-heavy,” with heavier segments (thigh, pelvis, and trunk) supported by lighter lower limbs. Consequently, the body center of mass moves at approximately 1 m (in adults) above the base of support. Thus, even small deviations of the center of mass position can result in substantial gravitational moments that accelerate the center of mass out of the base of support. Human walking is subject to internal perturbations resulting from the random variance in firing behavior of sensory afferents and motor neurons, as well as the performance of simultaneous motor tasks, such as breathing or more complex tasks like talking or scanning the environment. In addition, external perturbations, e.g. due to walking over uneven terrain or exposure to gusts of wind, may occur. Both internal and external perturbations can cause deviations in center of mass position with respect to the base of support. Given that walking is inherently unstable, ongoing (intermittent or continuous) stabilization is needed, to avoid cumulative effects of small perturbations over time. When major perturbations occur, which are here defined as external mechanical events that disturb the relation between center of mass and base of support beyond the variance observed in unperturbed gait, the need for stabilizing control is evident.

Stabilization of gait can be achieved by passive and active mechanisms. Passive stabilization depends on the passive mechanical properties (stiffness, damping and inertia of the human body), whereas active mechanisms involve modulation of neural drive in response to sensory information resulting in modulations in muscle activity. Passive mechanisms require no control effort and may have no direct energetic costs associated with them, but may not be amenable to change. Active mechanisms are presumably more adaptive to task requirements and may be more amenable to improvement by training in the long term. Active and passive mechanisms are probably used in parallel. A simple two-dimensional (sagittal plane) model of a bipedal walker can be stable without any form of active control. In such a model, the forward fall of the center of mass is arrested on a step-by-step basis through adequate foot placement resulting from the model’s passive dynamics (Mcgeer, 1990). The ground contact force created by the new stance leg after foot placement passes in front of the body center of mass and thus creates a backward moment with respect to the center of mass, which brakes the forward fall. However, these passive models cannot deal with perturbations of realistic magnitudes and three-dimensional model versions are unstable in the mediolateral direction (Kuo, 1999). This indicates that additional active control must be exerted to horizontally accelerate the center of mass in the desired direction, whenever errors in control or external perturbations cause deviations in the center of mass from its planned trajectory. Active control can result in changes in the walking motion to anticipate the effect of perturbations (feedforward control) or changes in muscle activity and hence movement in reaction to perturbations (feedback control). Feedback control requires information on the state of the system, that is, on the current center of mass state and its relation to the base of support.

We assume that feedback of the center of mass state is used to stabilize gait. Muscle activity after a perturbation cannot be explained by local (joint level) feedback alone (Safavynia & Ting, 2013), suggesting that responses are based on integrated information, possibly contributing to a center of mass state estimate. This information arises from multiple sensory modalities, including visual (Reimann et al., 2019), vestibular (Magnani et al., 2021; Reimann et al., 2017), and proprioceptive (Arvin et al., 2018; Roden-Reynolds et al., 2015) systems. Manipulations of each of these modalities during walking elicits responses that would be expected to “correct” the illusory movement caused by these manipulations. Note that it is not known whether an explicit estimate of the center of mass state is indeed generated and used to stabilize gait, but a proxy of this information seems required for effective feedback.

In this paper, we present methods to assess the feedback component in the control of human walking at three levels: 1) the level of center of mass movement and horizontal ground reaction forces, 2) the level of center of mass movement and foot placement (the dominant mechanism to control horizontal ground reaction forces) and 3) the level of center of mass movement, muscle activity and the joint moments. The methods presented can hopefully be of use to assess control of gait in a continued effort to better understand how gait is stabilized and how aging and disease affect this process. In addition, we hope and expect that they can be translated into tools for clinical assessment, to diagnose problems in stabilization of gait and to monitor the effect of interventions to improve this. We specifically aim to show that center of mass state feedback can be used to analyze feedback control in unperturbed gait. This would enhance their applicability (e.g. for clinical screening, where applying perturbations may not be feasible).

## 2. CONTROL OF HORIZONTAL GROUND REACTION FORCES

If feedback is used to stabilize the center of mass relative to the base of support, it is likely that deviations of the center of mass state (position and velocity) form the planned state are sensed and that these trigger actions to correct the deviations. These actions would be reflected in accelerations of the center of mass relative to the base of support in the opposite direction. In the absence of changes in the base of support position (e.g. single stance phase), these accelerations are of course directly proportional to the horizontal ground reaction forces (Newton’s second law). Even with adjustments in foot placement altering the base of support position, horizontal forces are required to control medio-lateral velocities during steady-state walking. Thus, if feedback is used, a negative regression between the center of mass state and horizontal ground reaction forces at a later point in the gait cycle is to be expected (e.g., we assume that a deviation in center of mass velocity to the left at time *t* will result in an center of mass acceleration to the right at time *t + δ* to return to a steady-state motion). Does such a correlation exist? We present an answer here that has not been published previously, so we will provide a detailed overview of the methods used, and results obtained.

### 2.1. METHODS

#### 2.1.1. FEEDBACK MODEL

To analyze correlations between center of mass state and horizontal ground reaction forces, it is useful to introduce the extrapolated center of mass (*xCoM*). Hof et al. (2005) showed that with some simplifying assumptions (modeling the human as a linearised inverted pendulum), the control of the center of mass state can be simplified into the control of a single variable, the *xCoM*. The *xCoM* is a weighted sum of the horizontal position and velocity of the center of mass:

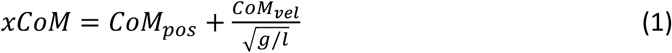

with *CoM*_*pos*_ = center of mass position, *g* = the magnitude of the gravitational acceleration and *l* = is the length of the inverted pendulum (approximated as 1.24 – 1.34 times trochanteric height (Hof et al., 2005)) and *CoM*_*vel*_ denoting the first-order time derivative (center of mass velocity). Using this simplification allows us to approach the problem at hand as a univariate regression between *xCoM* and the horizontal ground reaction force (*Fgr*) in the corresponding direction. We note that using the extrapolated center of mass is not necessary, a bivariate regression analysis using *CoM*_*pos*_ and *CoM*_*vel*_ as predictors is possible and may be preferable when the kinematics deviate more from inverted pendulum mechanics, e.g., in running or perturbed walking.

Our aim is thus to model deviations in *Fgr* in either the mediolateral or anteroposterior direction as a function of deviations of the *xCoM* that occurred earlier in time at an interval that would allow sensory information to be transferred to the central nervous system, to be processed, and to generate the appropriate motor responses. For simplicity we will assume a linear function. The modeling procedure is illustrated in Figure 1 for mediolateral *xCoM* and *Fgr*, but the same approach applies for anteroposterior control.

**Figure 1.**
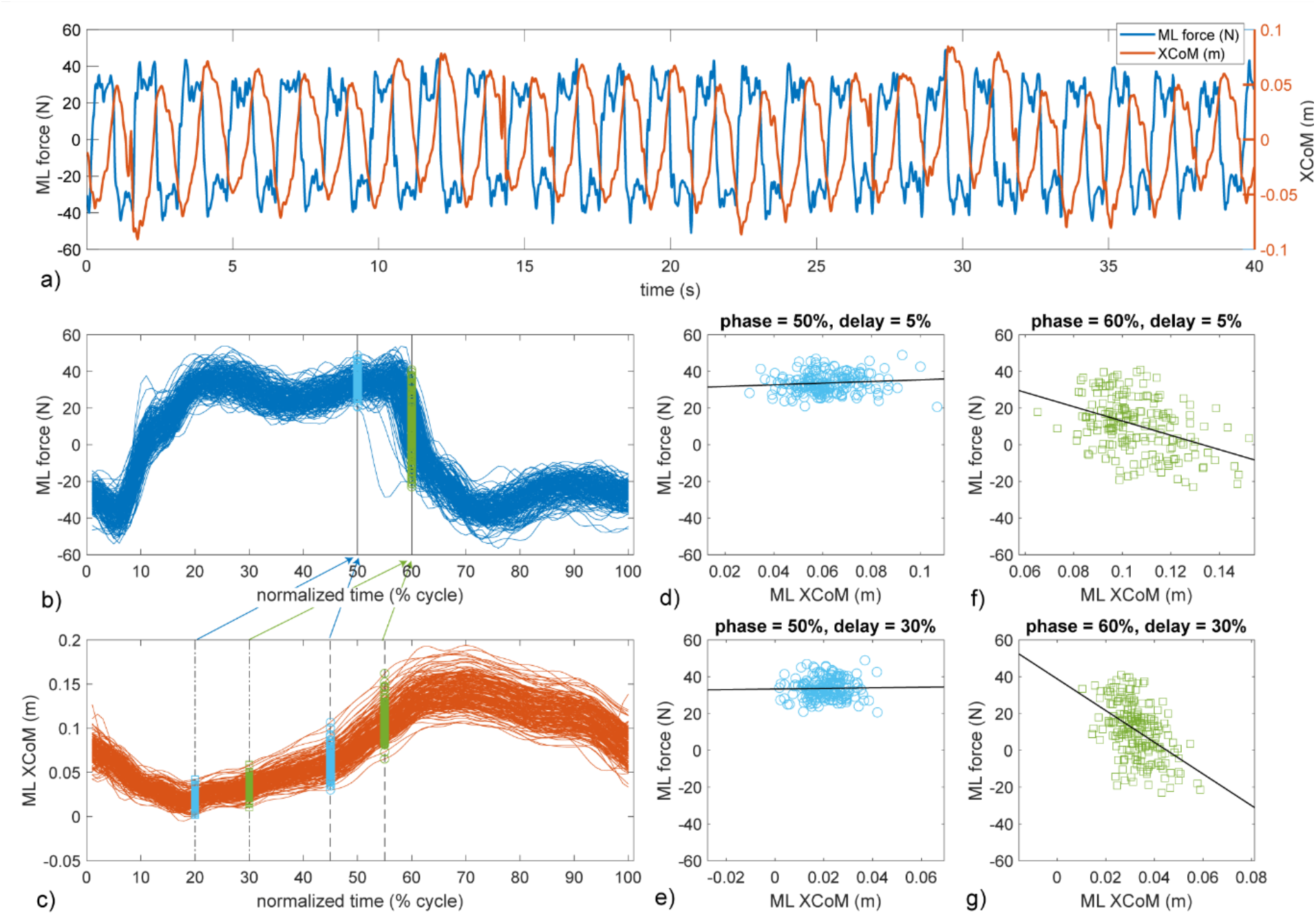
a) Time series of the mediolateral ground reaction force (Fgr on left y-axis) and extrapolated center of mass potion (xCoM on right y-axis) during walking of a healthy participant. Forty seconds of a trial of in total 5 minutes are shown. b) 200 normalized time series from left heel strike to left heel strike of Fgr extracted from the same time series. The light blue and green symbols refer to the data fitted by the regression models in panels d-g. c) idem for the xCoM relative to the stance foot. d) Scatter plot and regression line illustrating the prediction of Fgr at 50% of the gait cycle (right heel strike) from xCoM (referenced to the left stance foot) 5% of the gait cycle earlier. e) idem for xCoM 30% of the gait cycle earlier. f) idem for Fgr at 60% of the gait cycle and xCoM 5% of the gait cycle earlier. g) idem for xCoM 30% of the gait cycle earlier.

We consider all variance in *Fgr* and *xCoM* as deviations from a planned state, and thus assume the average gait cycle to reflect this planned state. We also assume human gait to be periodical, which allows us to divide time series of *Fgr* and *xCoM* (Figure 1a) into strides that can be time-normalized. Normalizing data from heel strike to ipsilateral heel strike to 100 samples yields an n x 100 matrix of *Fgr* and *xCoM*, with n the number of strides assessed (Figures 1b-c). In each stride, we re-reference *xCoM* data to the positions of the initial stance foot, which we will denote as *xCoM’*. We can now assess regression models of the following nature:

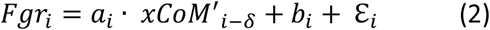

with *i* the % of the gait cycle and *δ* the delay (in % of the gait cycle) after which the feedback response becomes apparent, *a*_*i*_ and *b*_*i*,_ regression coefficients and *ε* _*i*,_ the residual error. This model can be fit to the n observations (strides) at each phase *i* (Figures 1d-g). Feedback responses are likely phase dependent (e.g.,Berger et al., 1984; Bruijn et al., 2015; Golyski et al., 2021; Magnani et al., 2021), so we assess this model separately for every phase *i ≥* 50%. We start at 50% of the stride to allow for using *xCoM* data during the preceding half of the gait cycle. If strides are normalized based on left heel strikes, the results will reflect how actions from right heel strike until left heel strike are determined by the center of mass state during the preceding phases beween left stance and right heel strike. Obviously normalizing strides from right heel strike to right heel strike will provide complementary, but assuming symmetrical gait, similar information. The value for the time delay, *δ*, is unknown and may vary between individuals and gait speeds, so we optimize *δ* across all phases in the gait cycles of a trial. This optimization is done based on maximizing the goodness of fit of equation 2. To find the best fit, we fitted the model for all possible values of *δ* and finding the value that yields the largest averaged negative correlation between *Fgr* and *xCoM* (Figure 2). Since feedback has to be negative to correct the center of mass towards the planned state, all positive correlations are discarded, negative correlations are Fisher transformed, that is the hyperbolic arctangent was calculated to obtain a normal distribution, averaged for each delay, and then inverse transformed.

**Figure 2.**
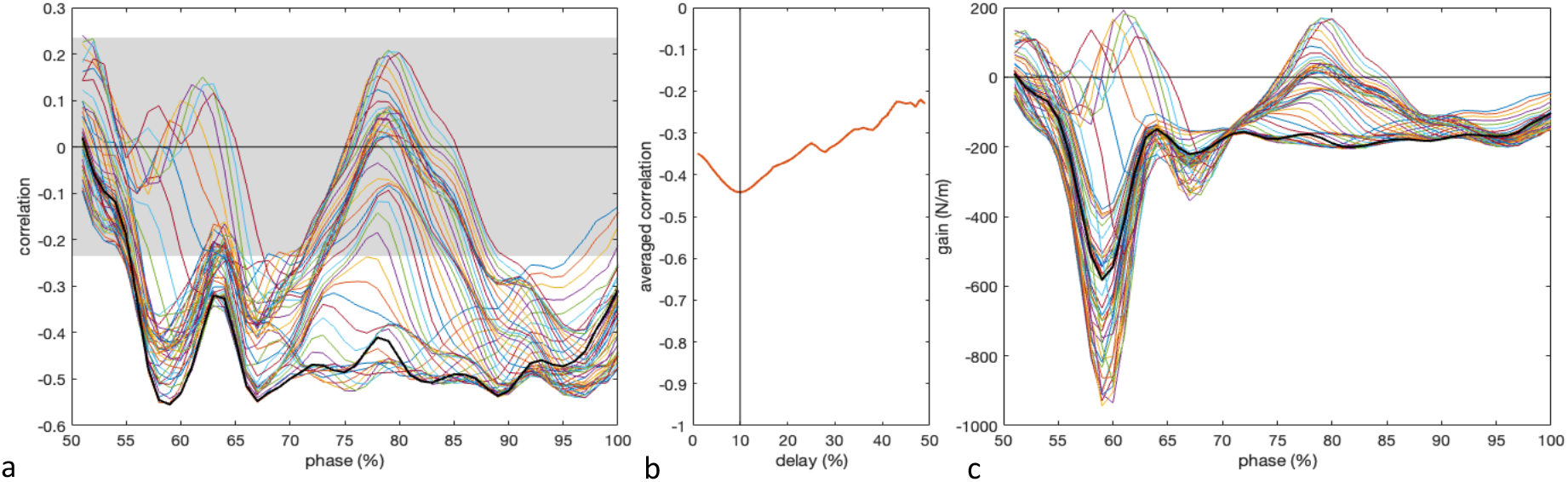
a) Typical example of the correlation between the mediolateral extrapolated center of mass (xCoM) and ground reaction force (Fgr) during the left stance phase with varying time delays between xCoM and Fgr (colored lines). The gray area indicates the interval within which correlations for the 200 observations (strides) used are non-significant. b) Averaged negative correlations and peak negative correlations as a function of the time delay between xCoM and Fgr. The black vertical line marks the delay with largest averaged negative correlation (10% of the gait cycle). c) Corresponding gains (regression coefficient a_i_ in equation 2) from the mediolateral extrapolated center of mass (xCoM) to the ground reaction (Fgr) with varying time delays between xCoM and Fgr (colored lines). The black line represents the gain obtained for the fit with the largest averaged negative correlation (10% of the gait cycle).

#### 2.1.2. EXPERIMENTAL DATA

Mediolateral ground reaction forces and whole-body center of mass trajectories during treadmill walking were obtained from healthy young adults, who participated in a previously reported experiment (van Leeuwen et al., 2020). Data of 30 participants were collected, but only data from 21 participants (14 females, 7 males; 31 ± 8 yrs, 70 ± 14 kg, 1.71 ± 0.08 m; mean ± SD), for whom a complete data set for the selected speed was available, were included in the analysis.

Participants were invited to walk on a treadmill at a normal (1.25 × √(leg length) m/s) normalized walking speed (Hof, 1996). In the experiment, stride frequency was controlled by means of a metronome, at a frequency set to the average preferred stride frequency determined during the final 100 steps of a familiarization trial. Trials lasted five minutes, to ensure that all trials contained at least 200 consecutive strides.

Participants walked on an instrumented dual-belt treadmill (Motek-Forcelink, Amsterdam, Netherlands). Ground reaction forces and moments were recorded from the force plates embedded in the treadmill and sampled at 200 Hz. Full body kinematics were measured using two Optotrak cameras (Northern Digital Inc, Waterloo Ontario, Canada) and sampled at 50 samples/s. Cluster markers were attached to the feet, shanks, thighs, pelvis, trunk, upper arms and forearms. Corresponding anatomical landmarks were digitized using a six-marker probe. We registered the location of several anatomical landmarks with respect to the cluster markers by pointing to the anatomical landmarks using a probe with six LED markers.

For all subjects, we analyzed 200 consecutive strides. Gait events (heel strikes & toe-offs) were detected based on the combined center of pressure as derived from force plate data (Roerdink et al., 2008). We estimated the mass of each segment using its length and circumference as predictors and regression coefficients based on gender (Zatsiorsky et al., 1990). The segment’s center of mass location was estimated to be at a percentage of the longitudinal axis of the segment (de Leva, 1996; Zatsiorsky et al., 1990). The full body center of mass was derived from a weighted sum over the body segments and its numerical derivative was used as an estimate of center of mass velocity. Both time series were combined to estimate the extrapolated center of mass trajectory as described above.

### 2.2. RESULTS

Figure 2 shows a typical example of the correlation between mediolateral ground reaction force and the extrapolated center of mass with varying time delays. Significant negative correlations are visible between approximately 55-75% of the gait cycle. Results for strides from right heel strike to right heel strike and for the anteroposterior direction were very similar. This result indicates that the horizontal ground reaction forces and hence the center of mass accelerations during the first half of single stance can, in this participant, be predicted from the extrapolated center of mass in the preceding swing phase. As can be seen in Figure 2b, the choice for the delay, does not strongly affect the result. The corresponding regression coefficients (*a*_*i*_ in equation 2), which can be interpreted as the gains of the feedback from the extrapolated center of mass to the force, are illustrated in Figure 2c.

The gain peaked around 60% of the gait cycle, which is the beginning of single stance (Figure 2c). The negative value of this peak indicates that deviations in the xCoM position during the left stance phase are corrected during the right stance phase. The pattern of the change in gain over the gait cycle is not strongly dependent on the choice of the delay between *XCoM* and *Fgr*.

The correlations and gains for all participants are shown in Figure 3 for the mediolateral and anteroposterior directions. The results presented above were clearly representative for the whole sample of participants analyzed. The delay estimates for the extrapolated center of mass to ground reaction force feedback are summarized in Table 1.

**Figure 3.**
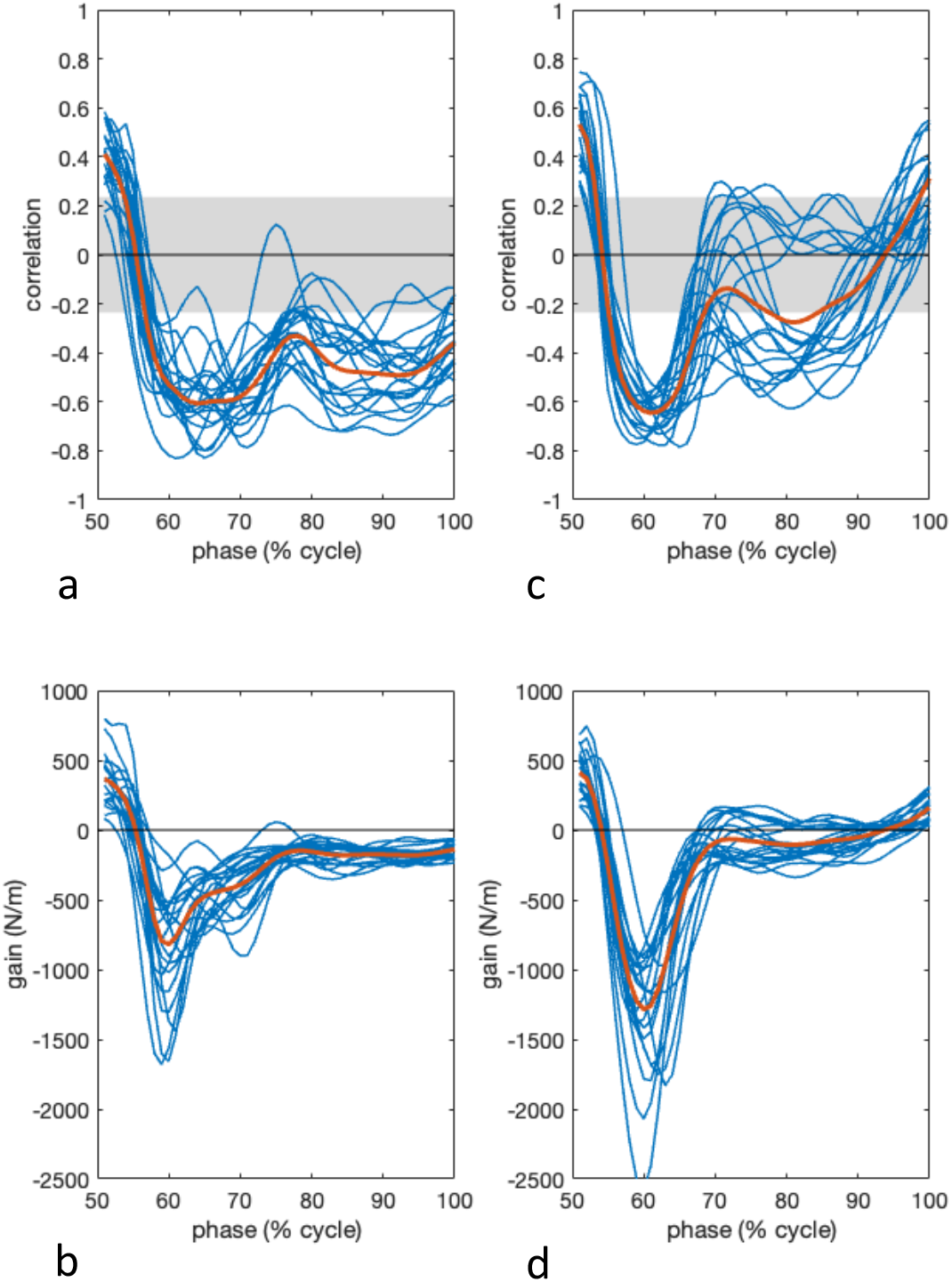
a) Correlations between the mediolateral extrapolated center of mass (xCoM) and ground reaction force (Fgr) during the right stance phase with selected time delays between xCoM and Fgr (blue lines). The black line represents the averaged correlation. The gray area indicates the interval within which correlations for the 200 observations (strides) used are not significantly different from 0. b) Corresponding gains from the mediolateral extrapolated center of mass (xCoM) to the ground reaction (Fgr) between xCoM and Fgr with ‘optimized’ time delays for all participants (blue lines). The black line represents the averaged gains. c) as in panel a, for the anteroposterior direction. d) as in panel b for the anteroposterior direction.

**Table 1.** Averaged estimated delays in percentage of the gait cycle and in ms with standard deviations in parentheses for both directions and for gait cycles starting at left and right heel strikes.

|  | mediolateral |  | anteroposterior |  |
| --- | --- | --- | --- | --- |
|  | left heel strike | right heel strike | left heel strike | right heel strike |
| delay (% cycle time) | 29 (12) | 32 (7) | 19 (12) | 12 (6) |
| delay (ms) | 323 (132) | 351 (79) | 215 (141) | 127 (63) |

### 2.3. DISCUSSION

The modeling procedure as described in section 1.1.1 has been implemented in Matlab and Python (code available from https://github.com/VU-HMS/Tutorial-stabilizing-walking). The function requires the times series of the extrapolated center of mass and the foot positions and heel strike events for both feet as inputs. As illustrated above, it allows estimation of the strength of the feedback coupling between the extrapolated center of mass and the horizontal ground reaction force, in terms of the correlation coefficient. In addition, it allows an estimation of the feedback gain and feedback delay. The strength of the feedback coupling and the gain are both clearly phase dependent as illustrated in Figures 2 and 3. The estimates for the feedback delay exceed values expected based on reflex delays, presumably because generating the required ground reaction forces may require placement of the swing foot (cf.Hof et al., 2010; Reimann et al., 2017; Wang & Srinivasan, 2014). The limited effect of the delay on the average (and peak) correlation (Figures 2a and b) may imply that information on center of mass movement is integrated over time to generate a more accurate response. The low variance of the averaged negative correlation as a function of the delay (Figure 2b) also renders the delay estimate less reliable and hence the marked difference in delays of anteroposterior force feedback between left and right stance phases may be an error. We also note that these estimates showed high between-participant variability.

## 3. CONTROL OF FOOT PLACEMENT

As argued previously (Bruijn & van Dieën, 2018; van Leeuwen, Bruijn, et al., 2022), foot placement (i.e. choosing the position of the leading stance foot) may be the main mechanism via which the horizontal ground reaction forces can be modulated. In this section, we will show how the control of foot placement can be estimated from kinematics of the center of mass and the feet. For this, we will follow the findings of Hurt (Hurt et al., 2010) and Wang & Srinivasan (Wang & Srinivasan, 2014), who showed that foot placement is linearly related to the center of mass state in the preceding step.

### 3.1. METHODS

#### 3.1.1. FEEDBACK MODEL

We define foot placement here as the position of the leading stance foot relative to trailing stance foot. We here focus on mediolateral foot placement, with positions medial to the trailing foot defined as positive and positions lateral to the trailing foot as negative. Note that in normal gait, foot placement is always positive. If foot placement (*FP*) is used to control the horizontal ground reaction force, and if this happens with respect to the center of mass state, there should be a relation between the deviations in center of mass state and deviations in subsequent foot placements. Assuming this relation to be linear (again for simplicity), the following should provide a good fit:

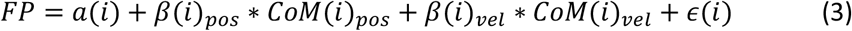

Identification of this model comes down to linear regression relating the dependent variable foot placement position (FP) to the independent variables center of mass position (*CoM*(*i*)_*pos*_) and velocity (*CoM*(*i*)_*vel*_), after which a residual variance (*ϵ*(*i*)) remains. We have implemented this procedure in our open source ‘Footplacement’ code(Bruijn & van Leeuwen, 2020), which is available in Matlab and Python format (https://github.com/VU-HMS/Tutorial-stabilizing-walking). This code requires as input the (AP or ML) center of mass (or a proxy thereof; some authors have used the pelvis position as such) trajectory, as well as the trajectories of both feet (we note that any marker on the foot can be used, and, under rigid body assumptions, should yield similar results. Here, we used the virtual calcaneus marker position), and the gait events (moments of heel strike, and toe off). To fit the model, we first divide the time series into steps, and time-normalize each step to 51 samples. Next, we define *FP* from the phase of the steps in which we are most sure that the foot does not move, i.e., the single stance phase.

Next, an important choice is the coordinate frame in which to express foot placement position and center of mass position. One could express these in the global coordinate frame, but this would lead to very high co-variance of both variables, as subjects may wander left and right (or back and forward) on a treadmill, which will be reflected in both the foot placement locations and the center of mass position. A more interesting choice may be to express both foot placement position and center of mass position in the coordinates of the previous stance foot, such that such variations in location are ignored^1^. In our code, selecting this option is achieved by setting the `removeorigin` parameter to 1. Another choice one could make now, is to decide to not be interested in the step width(length) per se, but only in the variations in step width(length). Doing so can be done by removing the mean from all variables of interest (FP and time normalized CoM positions and velocities). This does of course not lead to changes in *ϵ*(*i*), but it does omit estimation of the intercept *a*(*i*). In our code, this is done by setting the `centerdata` parameter to 1. After this, for each normalized time point *i* in the step cycle, we fit the linear regression equation, to obtain estimates of *β*(*i*)_*pos*_ and *β*(*i*)_*vel*_ as well as the variance explained by the model, and, for each step, the magnitude of the remaining residual, *ϵ*(*i*). This root mean square of the latter has also been interpreted as a measure of foot placement error (Jin et al., 2023; van Leeuwen, van Dieën, et al., 2022).

The estimates of *β*(*i*)_*pos*_ and *β*(*i*)_*vel*_ can be regarded as feedback gains (Reimann & Bruijn, 2023), and, in line with this, the fit of the model can be seen as a measure of quality of control. However, here, one should keep in mind that the R^2^ value is a ratio between the explained and total variance. We may obtain a higher R^2^ value by either an increased total variance (with equal explained variance, implying increased errors), or by reducing the residual variance. Hence, it may be useful to also investigate the magnitude of the foot placement error, the root mean square of *ϵ*. We note here that this model does not explicitly contain a delay. The model is fitted for the state variables at each phase *i* during the swing phase, the output is the foot placement at the end of that swing phase. Thus, the time difference between the instant of heel strike and the phase *i* can be considered the delay. The goodness of fit tends to saturate around mid-swing, and this may be used as indication of the delay.

#### 3.1.2. EXPERIMENTAL DATA

We used the data presented in section 1.1.2., for which the R^2^ has already been calculated and reported in a previous study (van Leeuwen et al., 2020). Results presented here may vary slightly from the results in that paper due the fact that a different subset of subjects was used for the calculations. *CoM*_*pos*_ and *CoM*_*vel*_ were calculated as explained in section 1.1.2. We focus on the mediolateral foot placement here, but a similar analysis can be performed for antero-posterior foot placement (Jin et al., 2023).

### 3.2. RESULTS

Figure 4a shows the explained variance of the foot placement model over the gait cycle, and figure 4b shows the residual variance for each prediction. As can be seen from this figure, the center of mass state at ipsilateral toe off can already explain as much as 60 % of the variance in foot placement. This percentage increases to roughly 85 % around heel strike.

**Figure 4.**
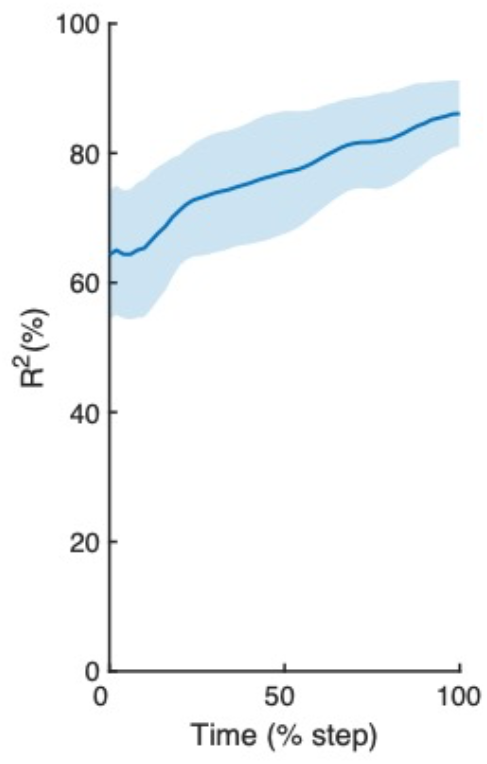
Percentage of variance in foot placement explained by preceding center of mass state, per percentage of the step-cycle (0% equals toe-off, 100% equals heelstrike). Already at the beginning of the step cycle, the center of mass state can explain about 60% of foot placement variance.

Figure 5 shows the regression coefficients *β*(*i*)_*pos*_ *and β*(*i*)_*vel*_. From this figure, it can be seen that the center of mass position plays a larger role in the prediction of foot placement, as *β*(*i*)_*pos*_ is approximately 5 times larger than *β*(*i*)_*vel*_. However, here one should keep in mind that deviations of *CoM*_*pos*_ and *CoM*_*vel*_ could have a different magnitude. Still, as this does not seem to be the case (See supplementary figure 1), deviations in center of mass position play a larger role in adjustments of foot placement (See also supplementary figure 2, which displays the normalized regression coefficients).

**Figure 5.**
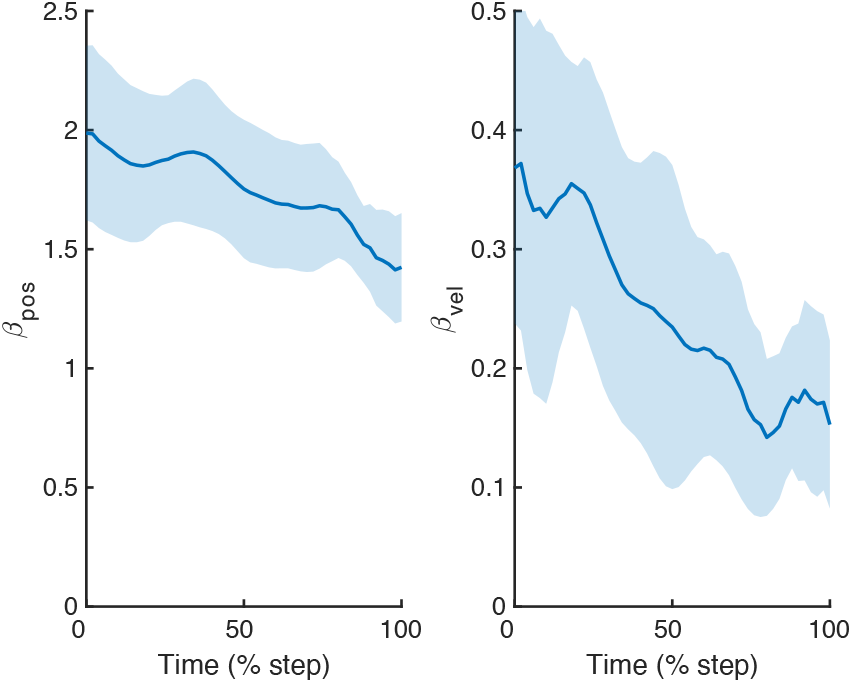
Regression coefficients β(i)_pos_ and β(i)_vel_ as a function of the phase of the step-cycle (0% equals toe-off, 100% equals heel strike). Note that foot placement is much more dependent on position of the center of mass than on velocity.

### 3.3. DISCUSSION

We have shown how, in a relatively easy way, one can estimate measures of foot placement control. These measures include feedback gains, as well as a measure of how well foot placement is coordinated to the center of mass state (R^2^). Of course, one may argue that the measures of foot placement control that we estimate might as well be measures of how passive dynamics influence the system, but this holds for most measures discussed in this tutorial, and will be discussed in the general discussion section.

Interestingly, the R^2^ has been shown to be lower in groups at an increased risk of falls, in older compared young adults and in stroke patients compared to healthy peers, suggesting that foot placement is impaired in these populations (Arvin et al., 2018; Stimpson et al., 2019). In addition, it has been shown that the R^2^ can be improved trough training (Hoogstad et al., 2022; Mahaki et al., 2023; Reimold et al., 2021; Reimold et al., 2020). Some of these studies have even explicitly used training based on the foot placement model, by deliberately improving (or degrading) participant’s foot placement accuracy (Heitkamp et al., 2019; Reimold et al., 2021; Reimold et al., 2020).

As mentioned in section 2.2.1, the residual variance of the model (*ϵ*((*i*)) has been interpreted as a measure of foot placement error. If one assumes that this is indeed an error in foot placement (and not simply some form of remaining noise), then it would seem logical that this error needs to be corrected, either using other mechanisms, or in the next foot placement. Indeed, van Leeuwen et al. (2021)showed that variance in mediolateral center of pressure shifts during single stance can be (partially) explained from the mediolateral residual variance. Similarly, Jin et al. (2023) showed that the variance in push of moment can be (partially) explained by the residual variance in antero-posterior foot placement. These findings together suggest that at least some of the residual variance in this foot placement model can be regarded as “foot placement error”. However, how much is actual “foot placement error”, and how much is “noise”, including measurement noise, remains open for future research.

*β*(*i*)_*pos*_ and *β*(*i*)_*vel*_ can be interpreted as feedback gains. A recent paper has shown that a linear inverted pendulum model can only show stable walking for a given range of such gains (Reimann & Bruijn, 2023). This suggest that it may be important to also consider the estimates for these gains. However, this manuscript also showed that humans typically have gains that are not within the range predicted by this model. The authors attributed this disagreement to the fact that the model lacked other stabilizing mechanisms (in particular ankle moment control). Still, difference in gains (albeit calculated in different ways) between healthy and patient populations have also been reported by Dean and Kautz (2015), highlighting the importance of studying these as outcome measures. The extrapolated center of mass, used as a descriptor of the center of mass state in the previous section of this paper, suggests that the ratio of these two gains should be 3:1 for position relative to velocity when calculated at foot placement (see also (Reimann & Bruijn, 2023)). This is different from the ratio of approximately five reported here, which may be due to differences in sensitivity to perturbations of these two state variables.

Here, like in most recent literature we have only included center of mass position and velocity in our foot placement models. We did so because we have found that in general, higher derivatives have limited additional explanatory value (Wang & Srinivasan, 2014). However, it should be mentioned that higher order derivatives of center of mass position can in principle be used, and earlier work by (Hurt et al., 2010) have actually included acceleration in their models. In our code, one can use derivatives up to acceleration, which can be done by setting the `order` variable; if set to 2, position and velocity will be included. When set to 3, acceleration will also be included.

## 4. CONTROL OF MUSCLE ACTIVITY AND JOINT MOMENTS

In addition to foot placement and other strategies, activating muscles in the stance leg to modulate joint moments can adjust the center of pressure under the stance foot to control the center of mass. This mechanism to control the center of mass appears to be used to correct for errors in foot placement during the subsequent stance phase (Jin et al., 2023), but is also used to counteract perturbations before foot placement can take effect, i.e., during the swing phase of the contralateral leg (Hof et al., 2010; Reimann et al., 2017). If activating muscles in the stance leg is used to stabilize the center of mass, the center of mass state should covary with delayed modulations in leg muscle activity (as measured by EMG) and joint moments of the stance leg. In this section, we will show how stance leg muscle activity and joint moments are related to the preceding center of mass state. For this, we will follow the findings of Welch and Ting (2008), who showed that reactive muscle activity is related to center of mass kinematics in perturbed standing and extend these findings to walking (published in (Afschrift et al., 2021)).

### 4.1. METHODS

#### 4.1.1. FEEDBACK MODEL

The relation between joint moment and center of mass kinematics during the stance phase is assessed using regression models of the following nature:

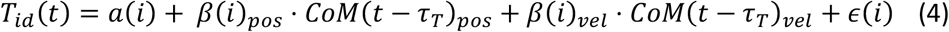

with *T*_*id*_*(t)* the joint moment, *β*_*pos*_ and *β*_*vel*_ position and velocity gains that potentially depend on the phase of the gait cycle (*i*), *CoM*_*pos*_ and *CoM*_*vel*_ the position and velocity of the whole-body center of mass with respect to the stance foot. *a(i)* depends on the phase in the gait cycle and represents the feedforward control. A neural delay of 100 ms (*τ*_*T*_) is used to model the time needed for signal transmission and sensory integration in the nervous system and electromechanical delay between muscle excitation and the development of force in the muscle (More & Donelan, 2018).

The correlation between muscle activity and center of mass kinematics during the stance phase is typically assessed using a slightly more complex regression model, in view of the fact that a muscle can generate a moment in only one rotational direction (e.g., soleus can generate a plantarflexion moment and the antagonistic tibialis anterior a dorsiflexion moment). We therefore introduce separate feedback gains for forward and backward directed deviations in COM trajectory with respect to a steady-state trajectory (Δ*COM*):

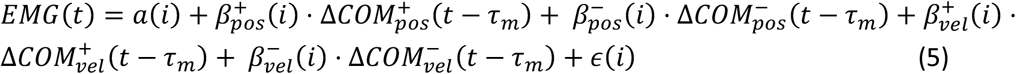

With *EMG(t)* the muscle activity measured with electromyography, 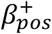 and 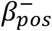 the gains for respectively the positive 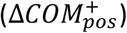 and negative 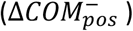 deviations in center of mass position and 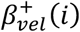 and 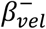 the gains for respectively the positive 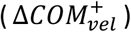 and negative 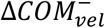 deviations in center of mass velocity. A neural delay of 60 ms (*τ*_*m*_) is used to model the time needed for signal transmission and sensory integration in the nervous system.

Fitting this model to experimental data is relatively straightforward and has been implemented in our open-source code (https://github.com/VU-HMS/Tutorial-stabilizing-walking). This code requires as inputs the center of mass position and velocity trajectories, the trajectory of both feet, joint moments and/or muscle activity and gait events. In contrast to the two previous methods, we do not express the center of mass motion as a function of the gait cycle, but search for every datapoint at time t (*EMG* or *T*_*id*_) the related center of mass kinematics at time *t-τ*. While for unperturbed walking the duration of a gait cycle is relatively invariant, this is not the case for perturbed walking with large variation in the duration of the stance phase and swing phase. Time normalizing the data would hence result in variable feedback delays between gait cycles, which is not desirable in this method because the time delay models specific processes (signal transmission, sensory integration and electromechanical delay).

With our open-source code one can fit a regression model on data pooled over a set of participants. It is important to nondimensionalize all quantities. Similar as in (Seethapathi & Srinivasan, 2019), we used center of mass height during quiet standing (*l*_*max*_), the gravitational acceleration (*g*) and body mass (*m*) to normalize positions, velocities and torques. Center of mass positions were normalized by *l*_*max*_, speeds by 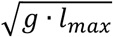, torques by *m*. *g*. *l*_*max*_. Normalization of EMG data is non-trivial (Besomi et al., 2020), hence we typically fit the second regression equation only to data of individual participants.

#### 4.1.2. DATASETS

We used the data presented in section 1.1.2. (unperturbed walking) and a dataset with push and pull perturbations applied at the pelvis at toe-off of the right leg (Vlutters et al., 2018). We evaluated in the unperturbed walking dataset the relation between center of mass state and ankle joint moments at multiple moments during the stance phase and in the perturbed walking data at 0.15 s after perturbation onset. The relation between center of mass state and ankle moments after anterior-posterior perturbations was already published (Afschrift et al., 2021) and we extended this analysis to the sub-talar, knee and hip joints and to medio-lateral perturbations.

### 4.2. RESULTS

Anterior-posterior center of mass position and velocity explained only a small portion of the variance in ankle plantarflexion moment during the stance phase in unperturbed walking (Figure 6b). The explained variance was highest during the first part of the stance phase (20-30% stance) and during push-off (Figure 6b). The low variance explained may be due to the relatively low signal-to-noise ratio in unperturbed walking. If this is indeed the case, then perturbed walking is expected to lead to stronger relationships.

**Figure 6.**
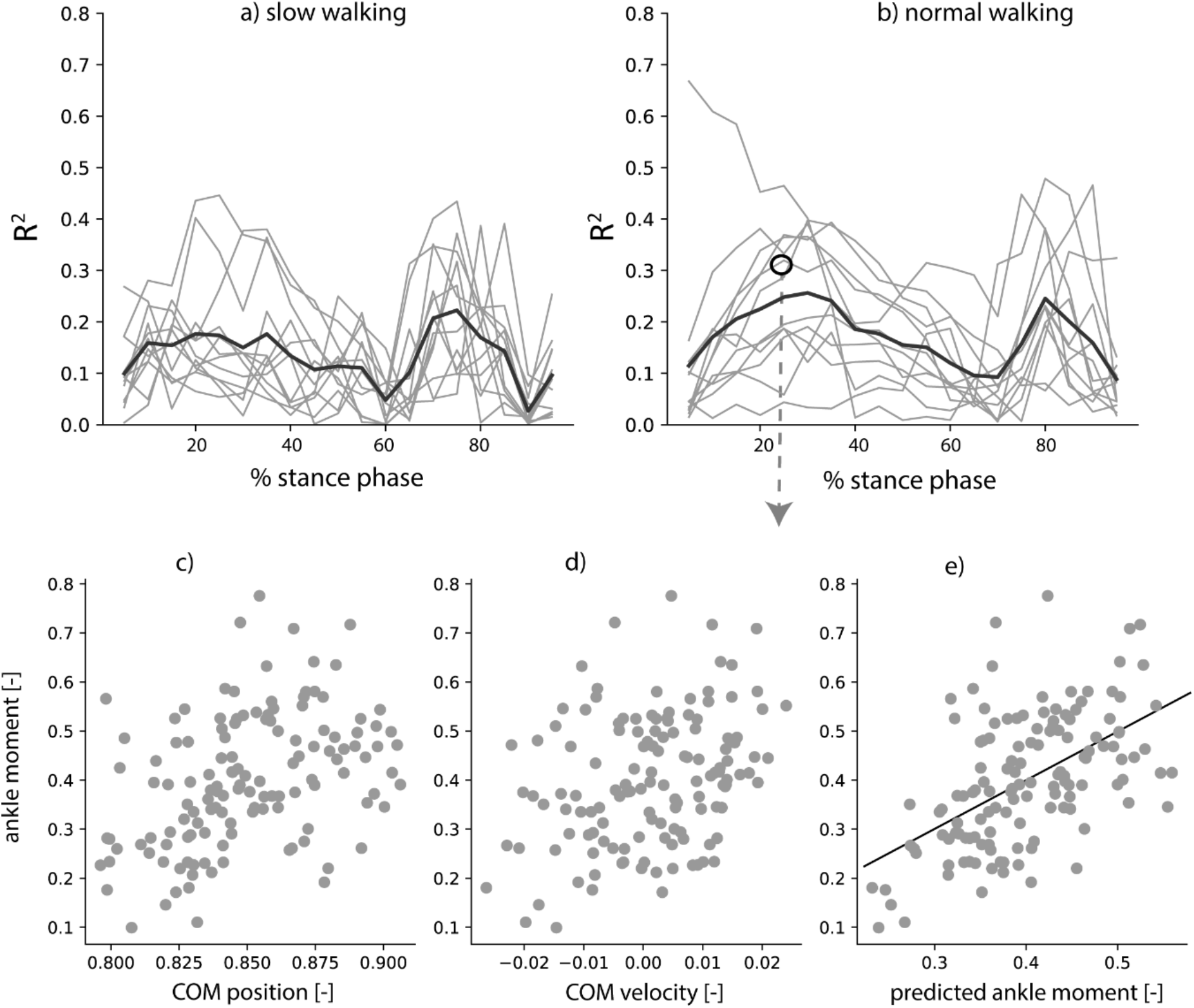
Variance in ankle joint moment explained by information on preceding center of mass kinematics as a function of the stance phase for slow (a) and normal (b) walking speeds. The relation between ankle moment and preceding center of mass position (c) and velocity (d) is shown for a representative example in normal walking (black dot in pane b).

As previously shown (Afschrift et al., 2021), we indeed found that anterior-posterior center of mass position and velocity can explain 86% of the variance in ankle moment after pull (posterior) and push (anterior) perturbations (Figure 7 A). The explained variance was highest for physiologically plausible delays (80-150 ms) and especially low for shorter feedback delays (see supplementary figure 3, which displays the explained variance as a function of the time delay). The relation between center of mass state and moments reflects an active control mechanism as tibialis anterior and soleus activity were also related to center of mass state (Figure 8 A). The variance explained by center of mass position and velocity feedback was lower for the subtalar joint moment (57%) knee joint moment (56%) and the hip joint moment (24%).

**Figure 7.**
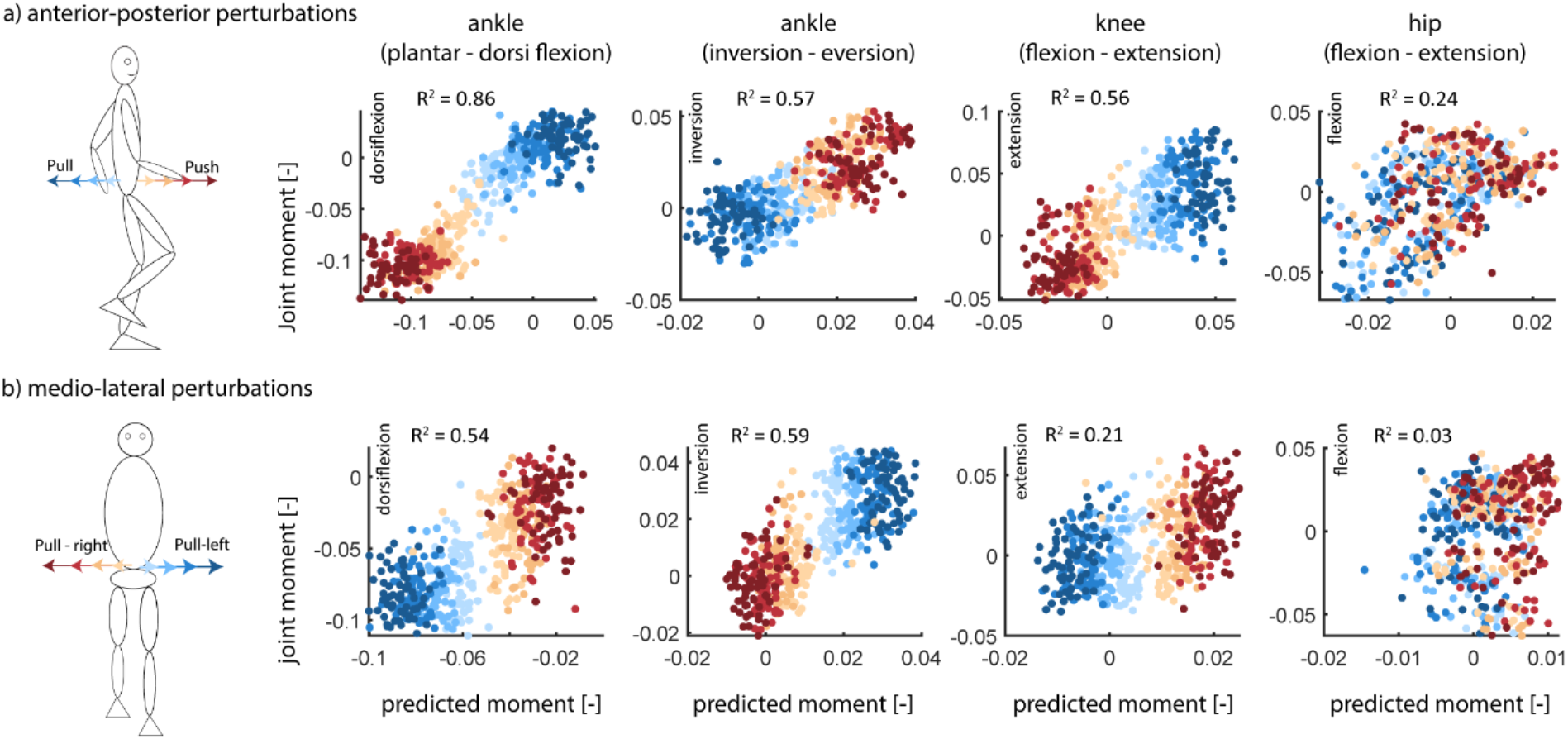
Relation between joint moment (ankle, subtalar, knee and hip) and predicted joint moment based on center of mass position and velocity for anterior-posterior directed perturbations (A) and medio-lateral directed perturbations (B) applied at the pelvis for the data pooled over all subjects.

**Figure 8.**
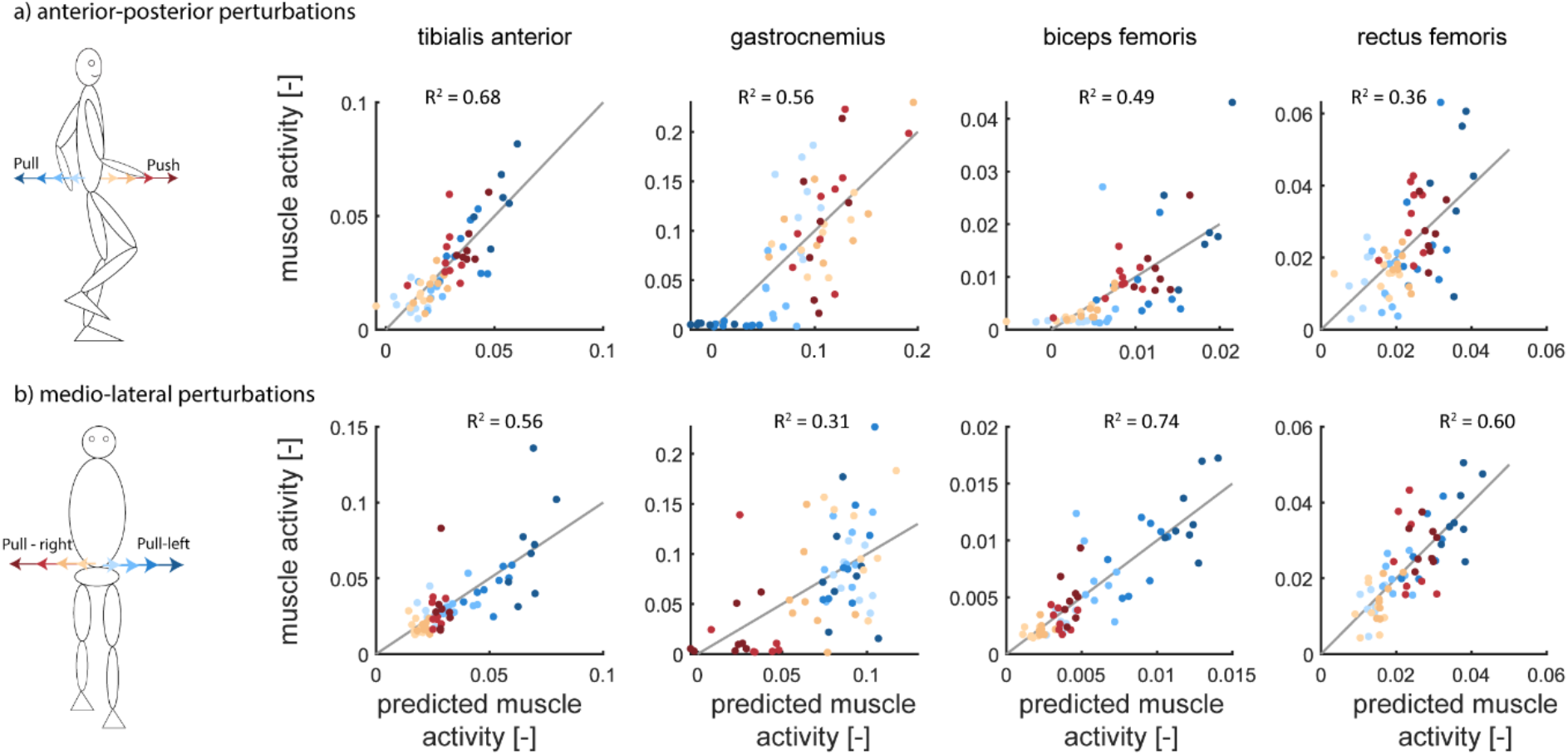
Relation between muscle activity measured with EMG (tibialis anterior, gastrocnemius, biceps femoris and rectus femoris) and muscle activity predicted from delayed center of mass position and velocity (regression model) for anterior-posterior (A) and medio-lateral directed perturbations (B) at the pelvis for a representative participant. Note that we did not perform this analysis on data pooled over all subjects due to the difficulty in normalizing EMG data.

In response to medio-lateral perturbations, the variance in ankle (54%), subtalar (59%), knee (21%) and hip (3%) joint moments explained by COM medio-lateral center of mass state was in general lower compared to the anterior-posterior directed perturbations. The related increases in subtalar inversion moment (Figure 6 B) and tibialis anterior activity (with an inversion moment arm, Figure 8 B) of the left stance foot in response to center of mass deviations to the left indicate that this is an active control mechanism.

### 4.3. DISCUSSION

We have shown that lower-limb joint moments and muscle activity are related to preceding center of mass position and velocity, particularly in perturbed walking, and to a lesser extent in unperturbed walking. This indicates that changing muscle activity in the stance leg, potentially in combination with other strategies such as foot placement and inertial strategies (e.g. hip strategy), is used to stabilize the center of mass during walking. The explained variance in ankle plantarflexion moment was much larger in perturbed walking (Figure 7) compared to unperturbed walking (Figure 6), which indicates that participants mainly used a foot placement strategy (Figure 4) in unperturbed walking.

During the balance recovery action after a perturbation, we found especially for the ankle plantar/dorsiflexion moment a strong relation with center of mass state. This is not surprising given that it is possible to modulate the center of pressure position under the stance foot directly with the ankle moment (so-called ankle strategy (Hof, 2007)). Furthermore, the potential movement of the center of pressure is larger in the sagittal plane than in the frontal plane due to the dimensions of the foot. It is therefore not surprising that the variance in plantarflexion moment explained after anterior-posterior perturbations was larger compared to that in the subtalar inversion/eversion moment after medio-lateral perturbations. While the variance explained in the ankle moment was largest, we also observed changes in knee and hip joint moments after perturbations, which could partially be explained by the center of mass state. This supports the idea that controlling balance by activating muscles in the stance foot is not a pure ankle strategy but is a full lower-limb mechanism.

## 5. GENERAL DISCUSSION

We have shown how in a relatively easy way, one can estimate measures of the control of ground reaction force, foot placement and joint moments during (perturbed) walking. These measures include feedback gains, as well as measures of “control quality”. Of course, one may argue that these estimates may reflect the passive dynamics of the system rather than active control (Patil et al., 2019). The relation between center of mass kinematics, ground reaction forces, foot placement and joint moments may reflect the passive dynamics of the swing leg and the (passive) intrinsic mechanical impedance of muscle fibers (i.e. preflexes). In addition, the models presented here shows correlations of center of mass state variables with horizontal ground reaction forces, foot placement, joint moments and muscle activity later in the gait cycle. However, of course this does not prove causation. Still, manipulations of sensory information, including, vestibular (Fettrow et al., 2019), visual (Reimann et al., 2019) and proprioceptive (Arvin et al., 2018) information show that foot placement, which is the major determinant of the horizontal ground reaction force, is adjusted in a way that is consistent with information that these sensory modalities would provide on center of mass state (van Leeuwen et al., 2020). Moreover, these adjustments are, at least in part, actively controlled as shown by associations with muscle activity (Rankin et al., 2014; van Leeuwen et al., 2020). In section 4, the relation between muscle activity (measured with EMG) and center of mass kinematics also clearly indicates that the participants actively responded to the perturbations, showing that the relation between joint moments and center of mass state is not just the result of the intrinsic mechanical impedance of muscles (e.g., force-length and force-velocity properties). The relative contribution of passive dynamics and active feedback control remains, however, unclear in this analysis. Accurate mechanistic models of swing leg dynamics and muscle contraction dynamics and methods to estimate muscle states are needed to unravel the relative importance of passive dynamics and active feedback control.

For the response to perturbations identified in section 4, it may not be a matter of debate that the models reflect feedback. For unperturbed walking (sections 2 and 3), one could suggest that the center of mass trajectory and foot placement both result from feedforward control, where both are planned in conjunction. However, the fact that illusory movements induced by sensory perturbations elicit predictable adjustments in foot placement (Arvin et al., 2018; Hendrik Reimann et al., 2018; H. Reimann et al., 2018) and ground reaction forces (Magnani et al., 2021) argues strongly in favor of an interpretation in terms of sensory feedback.

The methods described in sections 2 to 4, vary in terms of complexity of the instrumentation and analysis needed. While we used estimates of the center of mass state based on full-body kinematics in all sections, a single marker on the pelvis as a proxy of the center of mass may suffice for unperturbed gait (Yang & Pai, 2014). In section 2, ground reaction forces were measured on an instrumented treadmill, but these estimates could also be obtained from acceleration data, possibly derived from the same single marker or an accelerometer on the pelvis (and body mass). Potentially, inertial measurement units could replace optical motion capture, although reported results on the position of the feet relative to the center of mass (Refai et al., 2020) appear to be insufficiently accurate for the foot placement model of section 3. Methods based on such simpler instrumentation need to be validated in future studies, as this may be useful for wider application. In the context of future clinical application, it is also necessary to assess the reliability and sensitivity of the estimates obtained.

The models presented here assume feedback control back to a planned state of the center of mass, which is assumed to be reflected in the average state over multiple strides. This is a reasonable assumption for walking on a treadmill at a constant speed. Intentional changes of speed or heading direction would be reflected in changes of the center of mass state, as well as require changes in the control variables (ground reaction force, foot placement, joint moments), but since in these cases the planned state would be unknown, such modulations cannot be captured by the models presented here.

From a scientific perspective, important open questions remain about stabilizing feedback control of walking. While multiple sensory modalities contribute to the control processes described in this paper, the relative contribution of these sources of information is unknown. While it is likely that these contributions can be flexibly adapted to for example changing environmental conditions (Andreopoulou et al., 2015), a process called sensory weighting, this process has been studied mainly for static postural control and not for walking. Furthermore, it is not evident that this sensory information is used to obtain an actual estimate of the center of mass state or whether it provides an estimate of some proxy that is similar enough to the center of mass state to allow control of the center of mass. It is also unclear from our analysis if subjects adapted their walking motion to anticipate perturbations (feedforward control) and how these potential anticipatory changes in walking motion interacts with reactive feedback control. Finally, it remains an open question if models should also consider control of the body’s angular momentum in addition to- or maybe even instead of-the control of center of mass state (Pijnappels et al., 2004; van Mierlo et al., 2022). It is yet unclear how humans adjust joint moments and foot placements when there are conflicts between control of angular and linear momentum (e.g. adjusting foot placement to control linear and angular momentum after a trip perturbation). We hope and expect that the models presented here will support answering some of these questions.

## ACKNOWLEDGEMENTS

Sjoerd M. Bruijn was supported by a grant from the Netherlands Organization for Scientific Research (016.Vidi.178.014), https://www.nwo.nl/en/

## Footnotes

1 It should be noted that what we described here as “foot placement position” FP, has been dubbed step width by some authors. These authors have generally used the center of mass position to reference the foot placement to, so that foot placement is how far the foot is placed lateral to the center of mass (instead of the stance foot). This logically means that some of the variance that is present in both foot placement position and center of mass position is negated, and, as such, leads to lower values for explained variance.

