## Supplementary figures and images for "ASSESSMENT OF STABILIZING FEEDBACK CONTROL OF WALKING, A TUTORIAL"

### supplementary figure 1

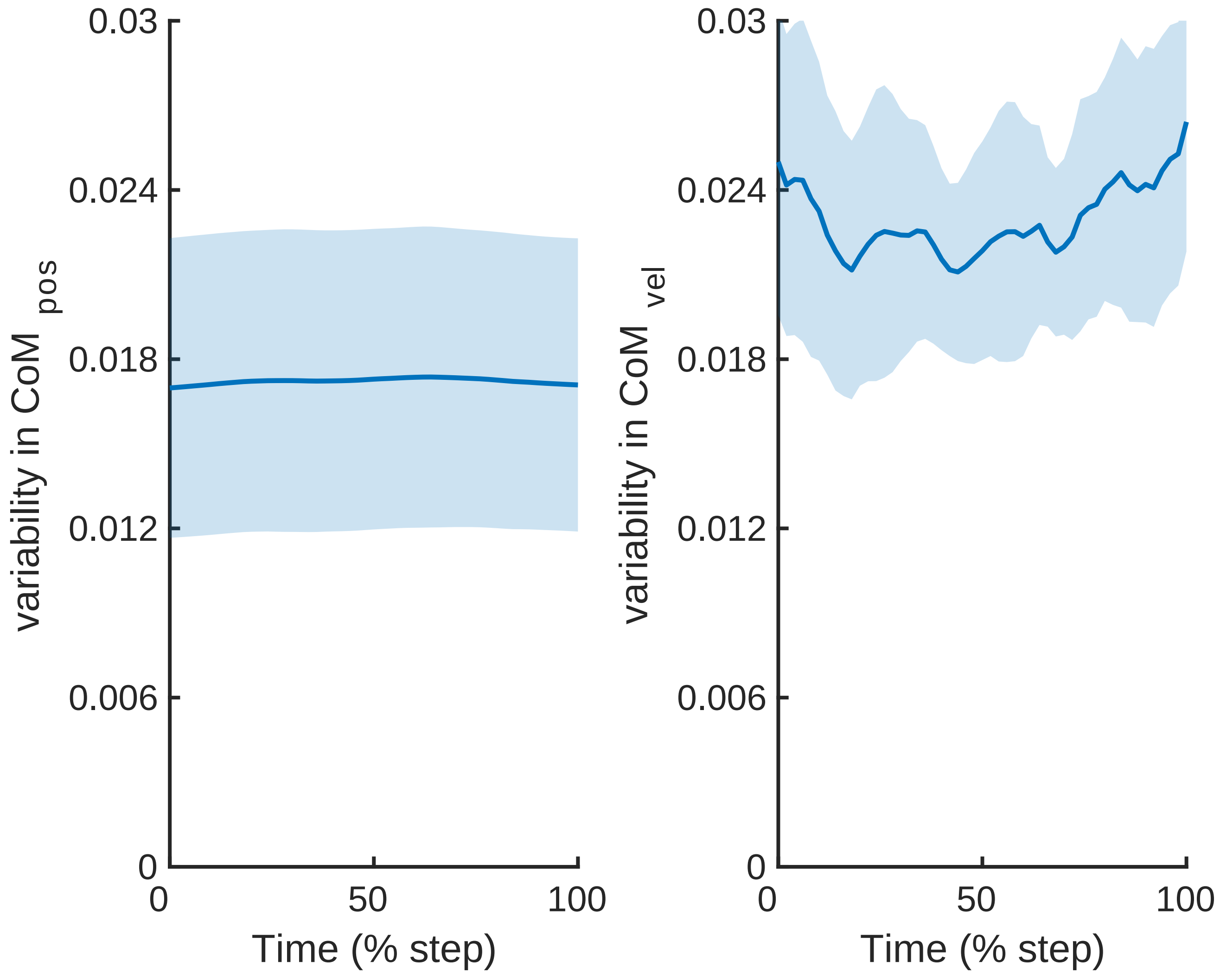

### supplementary figure 2

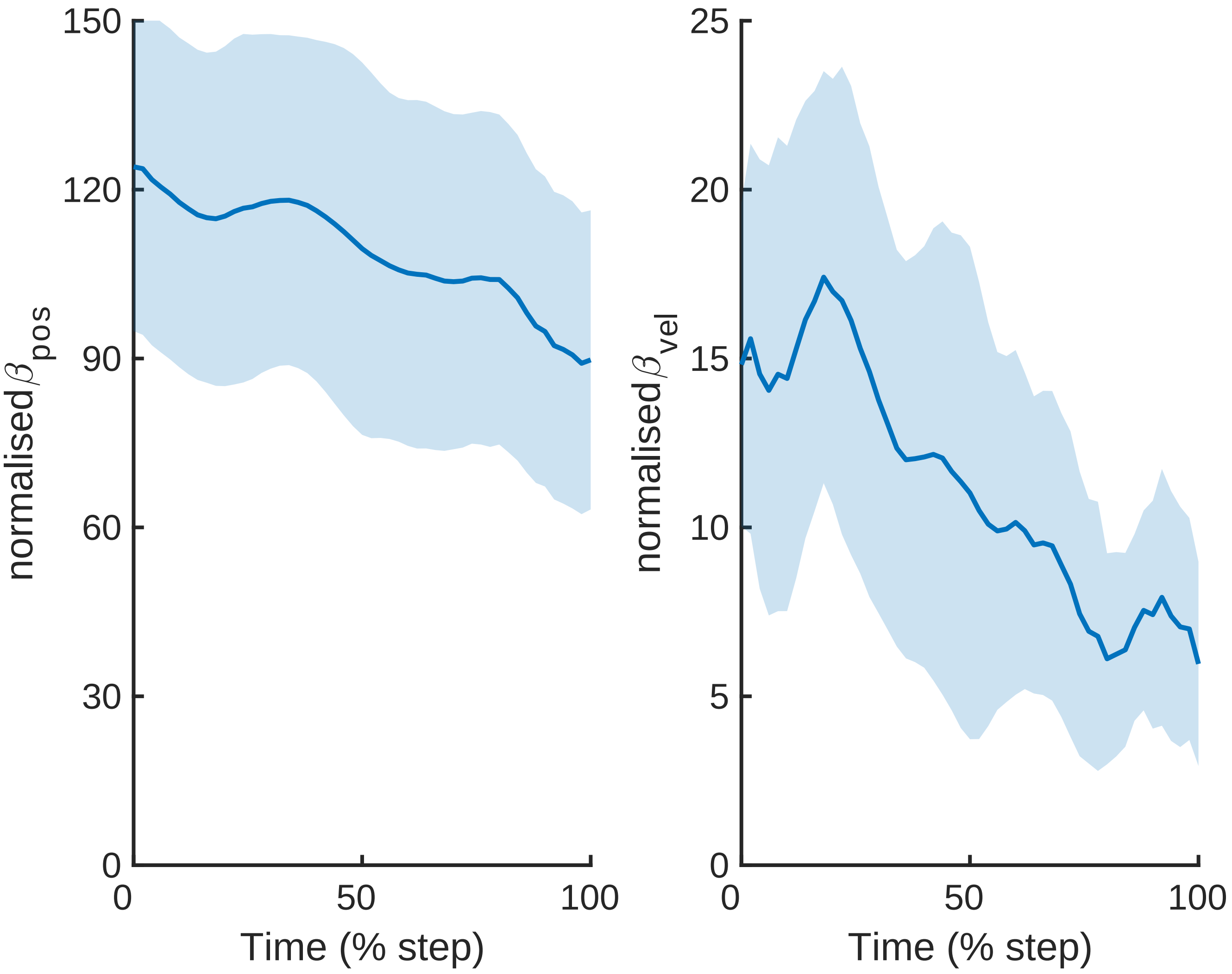

### supplementary figure 3

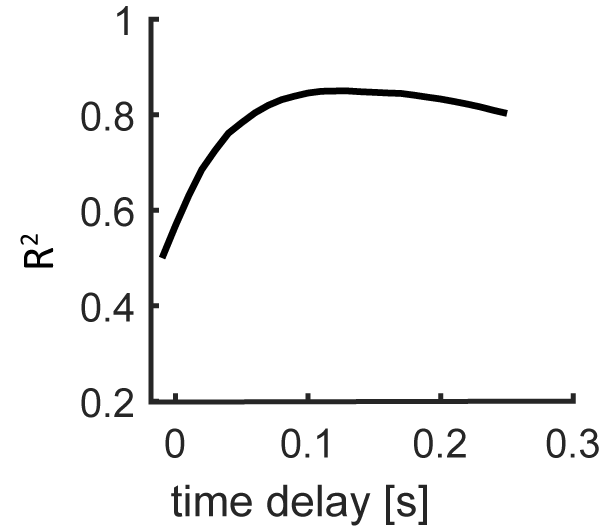
